# Unraveling the origin of Glucose mediated disparate proliferation dynamics of Cancer stem cells

**DOI:** 10.1101/2021.01.20.427381

**Authors:** Tagari Samanta, Sandip Kar

## Abstract

Cancer stem cells (CSCs) often switch on their self-renewal programming aggressively to cause a relapse of cancer. Intriguingly, glucose differentially triggers the proliferation propensities in CSCs in an origin-dependent manner by controlling the expression of the key transcription factor like Nanog. However, the factors that critically govern this glucose-stimulated proliferation dynamics of CSCs remains elusive. Herein, by proposing a mathematical model of glucose-mediated Nanog regulation in CSCs, we showed that the differential proliferation behavior of CSCs can be explained by considering the experimentally observed varied expression levels of key positive (STAT3) and negative (p53) regulators of Nanog. Our model reconciles various experimental observations and predicts ways to fine-tune the proliferation dynamics of specific CSCs in a context-dependent manner. In future, these modeling insights will be useful in developing improved therapeutic strategies to get rid of harmful CSCs.

## Introduction

Cancer is a disease related to uncontrolled cellular proliferation. It is known experimentally that a malignant tumor normally consists of different cell-types having a range of proliferation propensities (Reya et al., 2001; Yu, 2007). A subpopulation of cells within the cancerous tumor, however, unlike the normal proliferating cells, are capable of both self-renewal and can even differentiate into another cell-type depending on the external cues (Yu et al., 2012). These cells are termed cancer stem cells (CSCs), as they demonstrate stem cell-like properties (Kreso et al., 2014). Interestingly, the properties of differentiated CSCs are mostly like normal stem cell and their proliferation rates are much lower compared to an undifferentiated ones (Beck et al., 2013; Shackleton et al., 2009). On the other hand, the undifferentiated CSCs mainly generate tumors through abandoned self-renewal. The regrowth of such tumors from CSC needs nutrient acquisition like any other cell. It is observed that CSCs preferably use a glucose-intensive glycolysis pathway to get the required nutrients for extensive proliferation. Therefore, glucose is expected to have a significant role in survival as well as the proliferation maintenance of CSCs (Butler et al., 2013; Hirschhaeuser et al., 2011; Jang et al., 2013; Muñoz-Pinedo et al., 2012; Zhao et al., 2013). How glucose mediates these responses and orchestrates the proliferation pattern of CSCs, remains an important open-ended question.

In this context, Flavahan et al. measured the expression levels of various stem cell and differentiation markers for different glucose concentrations in CSCs known as human Brain Tumor Initiating Cells (BTICs) (W. A. Flavahan et al., 2013). They reported that in presence of restricted glucose concentration (2.5 mM), the expression levels of the stem cell markers (Nanog, Oct4, and Sox2) are quite high leading to a higher proliferation response of BTICs, while differentiation of these cells is non-existent. This observation is quite surprising, as according to the conventional ‘Warburg effect’ (Heiden et al., 2009), CSCs are expected to proliferate more prominently at high glucose concentration. The observation also contradicts the report by Derr et al., which shows shorter survival of patients with high glucose in brain tumors compared to the patients with low glucose concentration (Derr et al., 2009). Even for CSCs in other cancer types, such as pancreatic cancer, PANC-1; ovarian cancer, A2780; and glioblastoma, GS-Y03, one found higher rates of proliferation with increasing doses of glucose concentration. In this regard, Shibuya et al. (Shibuya et al., 2015) cultured human CSCs in presence of 5 mM glucose (physiological glucose concentration) and 26.2 mM glucose separately, and they observed that the CSCs preferentially differentiated at low glucose culture concentration. They have done both the pharmacological inhibition and genetic knockdown of glucose transporters and narrated that the inhibition of glucose transport restricts the self-renewal capacity of CSCs. This reveals that there is cell-type specificity within CSCs in relation to the proliferation pattern as a function of increasing levels of glucose. How this cell-type-specific proliferation is maintained in different CSCs needs to be unraveled properly to develop better therapies against uncontrolled proliferation in the future.

Nanog is one of the key molecular regulators of stem cell dynamics and this is true even for CSCs as well. The Nanog dynamics has been extensively studied experimentally to understand the proliferation as well as differentiation inclinations of CSCs. In the context of glucose-mediated proliferation of CSCs, Nanog has been monitored experimentally to understand whether CSCs opt for proliferation or undergo differentiation for specific cell-types. However, very little is known about the probable complex biological mechanism via which glucose affects the Nanog regulation and tilts the balance of proliferation to differentiation in CSCs. Thus, to understand the influence of certain interactions, mathematical modeling can be used as an important tool to explore the system at least qualitatively. In the literature, mathematical modeling has been employed extensively to study the proliferation and differentiation dynamics of stem cell regulation. Most of these studies reported qualitatively how the key stem cell markers induce self-renewal and differentiation under different biochemical scenarios for stem cells and improve our knowledge about the dynamical transition happening within a stem cell population. However, there is no such model proposed to date, which can explain the variation of proliferation and differentiation pattern of CSCs as a function of glucose, by keeping Nanog as a central regulator of stem cell dynamics.

Keeping this in mind, in this article, we proposed a mathematical model centered around Nanog regulation in CSCs to provide the probable explanation behind the differential dynamics of CSCs under low or high glucose concentrations. Our modeling study reveals that the disparate dynamics of CSCs under similar levels of glucose can be explained if Nanog shows a reverse mushroom kind bifurcation in these systems. Herein, we demonstrated that the differential behavior of Nanog in CSCs is mainly governed by the relative expression levels of the p53 and STAT3 (signal transducer and activator of transcription 3) proteins downstream to glucose in a cell-type dependent manner. In this regard, we put forward another parallel model of Nanog regulation, where the same Nanog network can give rise to different kind of bi-stable dynamics of Nanog for the same p53 and STAT3 levels as depicted earlier to explain the dissimilar responses of CSCs in presence of low and high glucose levels. Moreover, our model qualitatively reproduces several experimental observations related to the proliferation of CSCs of various cell-types in presence of glucose and makes further predictions to change the course of the proliferation of the CSCs by perturbing the Nanog network in a cell-type-specific manner. Thus, our model provides the rationale behind the effect of glucose on unwanted proliferation in a cell-type dependent fashion.

## The Model

In literature, several models with increasing complexities had already been proposed to study the Nanog dynamics within embryonic stem cells (Akberdin et al., 2018; Chickarmane et al., 2006; Chikarmane and Peterson, 2008). Here we proposed a minimalistic model of Nanog regulation (**Fig. 1**) based on the previous studies and improving it further using new information based on recent experimental literature. We subdivided the overall network model into two modules.

**Fig. 1.**
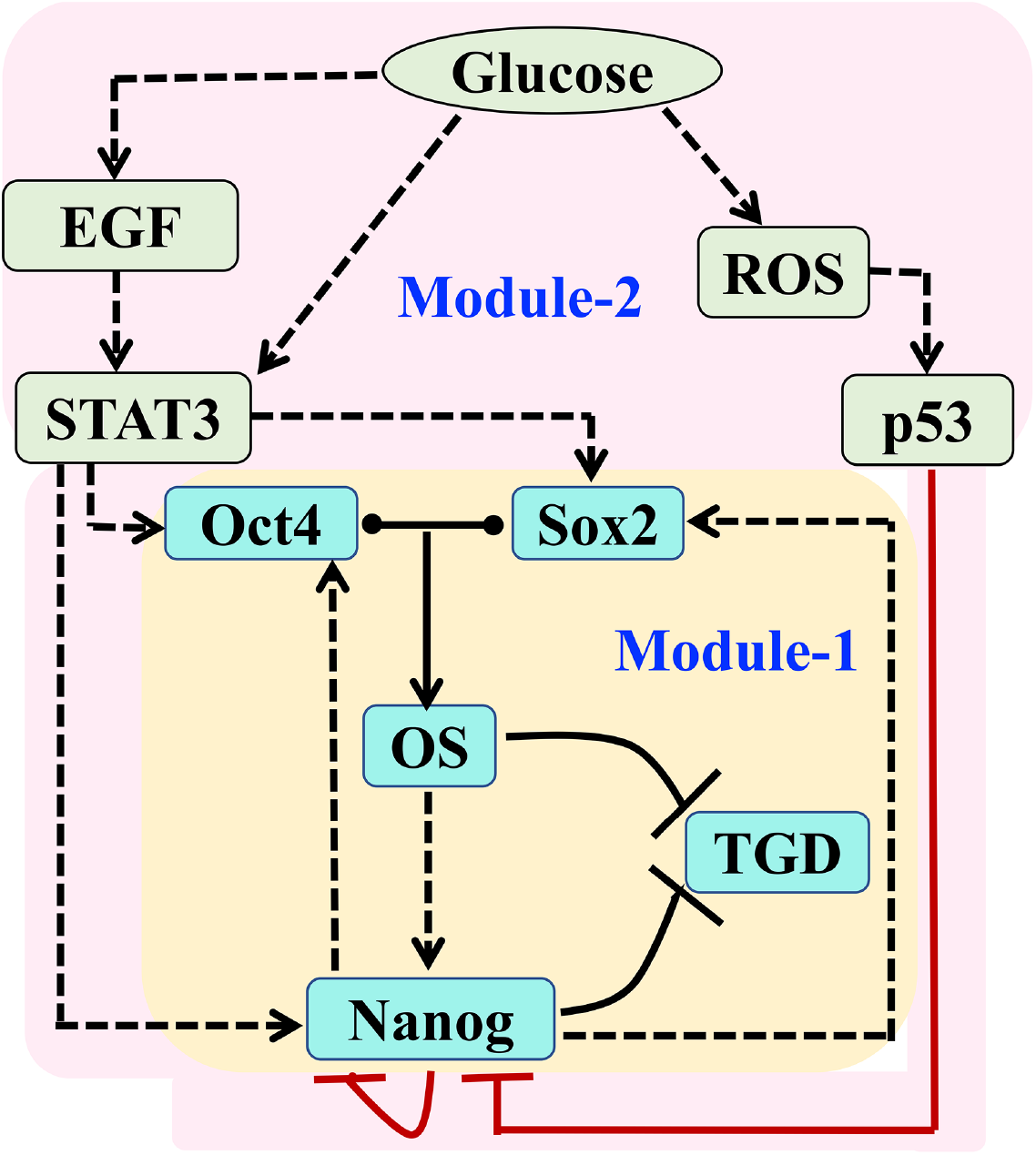
The Proposed Gene Regulatory Network of glucose-regulated Nanog dynamics in CSCs. The overall network is divided into two modules. **Module – 1** represents the core Nanog regulatory network of stem cell dynamics as depicted by Chikarmane et. al. (Chickarmane et al., 2006). **Module – 2** depicts how glucose regulates stem cell dynamics by altering Nanog through various positive and negative regulators. (The dotted and hammer-headed arrows represent activation and inhibition respectively. The arrow coming out from the middle of a dumbbell-shaped line indicates complexation-decomplexation events.)

### Module-1

To begin with, we considered an already proposed model by Chikarmane et al. (Chickarmane et al., 2006) that had successfully described the bi-stable Nanog dynamics operative within stem cells. In their model, they have considered an interplay between three core transcription factors of stem cells namely Nanog, Oct4, and Sox2 (**Module-1**, **Fig. 1**), and phenomenologically modeled the Nanog regulation in stem cells. According to that model, high expressions of these three transcription factors drive the stem cells towards proliferation, whereas lower expressions lead to differentiation. Following Chikarmane et al., we have considered that the heterodimeric complex of Oct4-Sox2 (OS) serves as an activator for all three transcription factors (Boyer et al., 2005; Festuccia et al., 2012; Ivanova et al., 2006; Loh et al., 2006). Furthermore, we assume that Nanog auto-represses its transcription (Navarro et al., 2012). In addition to this, we incorporated that there is a repressive effect of the Oct4/Sox2 (OS) complex on Nanog at high Oct4 concentration as indicated by the study of Karwacki-Neisius et al. (Karwacki-Neisius et al., 2013). The dynamical equations for the corresponding variables are provided in **Table-S1**, and the definition of the variables are given in **Table-S2**.

### Module-2

In **module-2**, we included some of the key regulators of Nanog that are activated in one way or another by glucose stimulation. In their work, Chickarmane et al. (Chickarmane et al., 2006) had speculated that upstream positive or negative signals might be present in the stem cells, and depending on the extent of such positive and/or negative regulations, the stem cell either undergoes proliferation or opts for differentiation. What are those positive or negative signals and how these signals are activating varied responses of Nanog are not mentioned? Herein, we tried to provide a more elaborate description of this part of the network under the influence of glucose, which ultimately dictates the disparate regulation of Nanog in CSCs. In literature, it is known that p53 negatively regulates Nanog expression post DNA damage and helps in the initiation of differentiation as well as inhibits self-renewal of stem cells (Lin et al., 2005; Zhao and Xu, 2010). Thus, p53 can be considered as a negative regulator of stem cell proliferation. Intriguingly, p53 activation gets regulated by glucose through reactive oxygen species (ROS) (such as hydrogen peroxide (H_2_O_2_), and superoxide (O_2_^−^), etc.) within stem cells. It is well-known that ROS signaling can induce DNA damage and stimulate p53 production (Ali Azouaou et al., 2015). In stem cells, glucose controls the production of ROS via several mechanisms, such as glucose autoxidation, activation of the reduced form of nicotinamide adenine dinucleotide phosphate oxidase, and production of advanced glycation end products (Cheng et al., 2016; Luo et al., 2018; Yu et al., 2011). Hence, glucose indirectly activates p53 activation via ROS. We included these interactions in our model using phenomenological terms (**Table-S1**).

We went further in the literature in search of some positive regulators of stem cell proliferation. We found that there are several positive regulators (Czerwinska et al., 2017; Du et al., 2016; Sato et al., 2012; X. Wang et al., 2017; Xu et al., 2017; Zhang et al., 2018), of stem cell proliferation dynamics, however, signal transducer and activator of transcription 3 (STAT3) (Du et al., 2016; Hadjimichael et al., 2017; Lv et al., 2017; Okumura et al., 2010; Zhang et al., 2017) and Epidermal growth factor Receptor (EGFR) signaling, (Di et al., 2013) seem to have a direct relationship with glucose stimulation. Epidermal growth factor (EGF) is known to promote stem cell proliferation via activation of STAT3 by phosphorylation and nuclear translocation (Saengboonmee et al., 2016). In fact, in presence of high glucose concentration, cell proliferation gets induced due to elevated EGF protein expression (Han et al., 2011). Experimental results even suggest that high glucose promotes cell proliferation via pathways apart from EGF, through direct activation of STAT3, which in turn positively regulates Nanog, Oct4, and Sox2 (H. Wang et al., 2017). Therefore, to study the effect of glucose on stem cells, we have considered both STAT3 and EGF as positive regulators of stem cells as the downstream target of glucose again by using phenomenological modeling terms (**Table-S1**).

We have translated the gene regulatory network shown in **Fig. 1** into an ordinary differential equation-based mathematical model by including both **module-1** and **module-2** (for details see **Supplementary information**). The parameters involved in the model along with their numerical values are provided in **Table-S3**. We have obtained some of the parameter values directly from the available experimental literature (shown in **Table-S3**). The other parameters are chosen in such a manner that the model qualitatively reconciles the experimental observation made for various types of CSCs under different glucose concentrations.

## Results and Discussions

### Model explains the glucose-mediated differential proliferative dynamics of CSCs

To begin with, we wanted to understand why CSCs from unlike origin behave differently as a function of increasing doses of glucose in terms of the dynamics of Nanog (stem cell marker) protein and a differentiation related protein (TGD) (**Fig. 2**). It is well-known that Nanog exhibits bi-stable dynamics in stem cells. Keeping this in mind, we performed a systematic bifurcation analysis of Nanog steady state dynamics as a function of glucose concentration. We found a bi-stable steady state dynamic of Nanog under the chosen parametric condition (**Table-S3**), where Nanog switches from a higher steady state (ON state) to a lower Nanog steady state (OFF state) to initiate the differentiation (**Fig. 2A**) with increasing doses of Glucose. The differentiation marker protein (TGD) shows just the opposite trend (**Fig. 2B**) to that of the Nanog dynamics. These results kind of resemble the brain tumor-initiating cells (BTICs), which preferentially undergo active proliferation (high Nanog and low differentiation protein) due to glucose deprivation (2.5 mM glucose concentration). However, for BTICs, differentiation (low Nanog and high differentiation protein) is promoted at higher glucose concentration (25 mM glucose) as shown by Flavahan et al. (W. A. Flavahan et al., 2013).

**Fig. 2.**
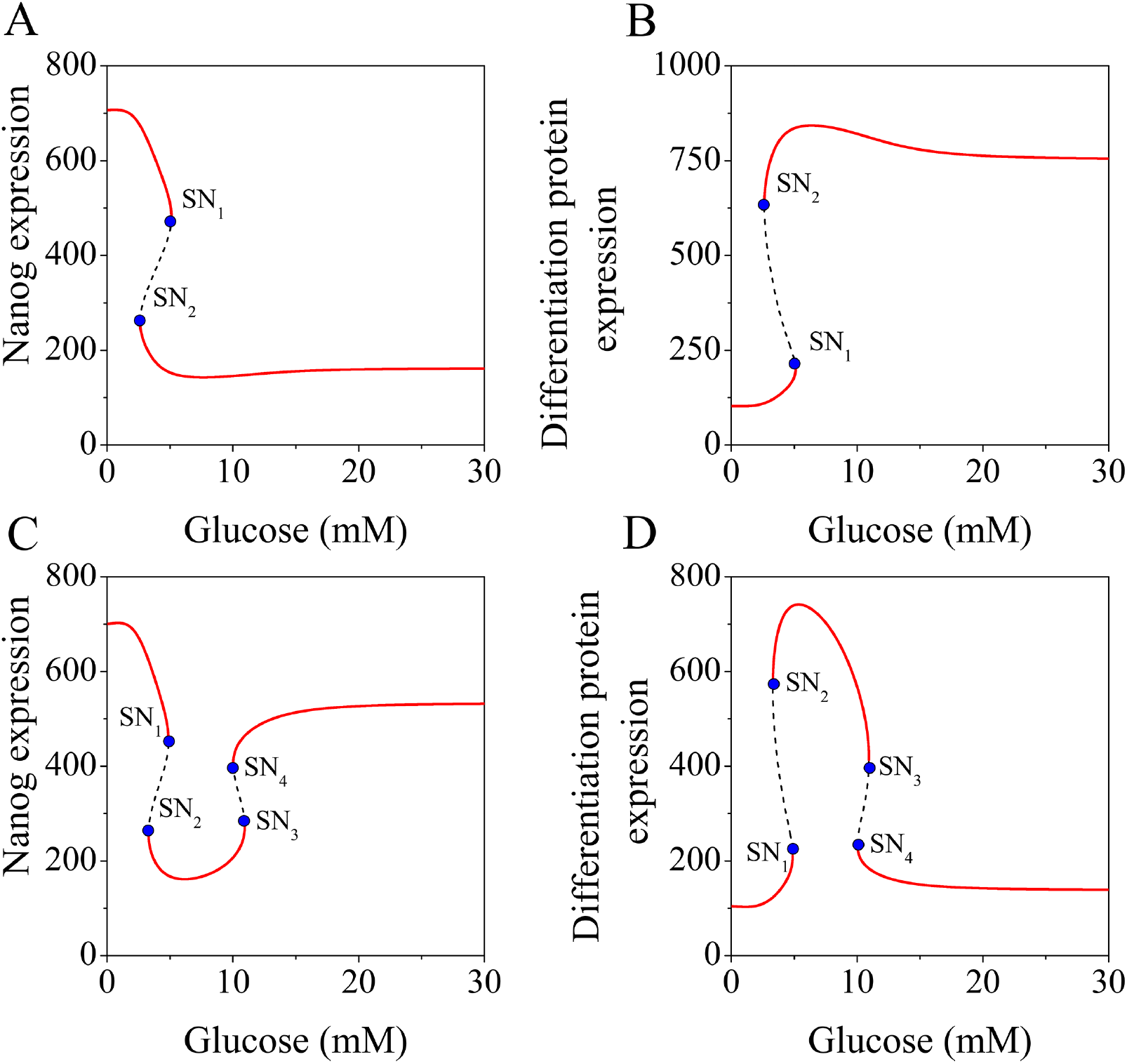
Bifurcation analysis of the model reveals the disparate proliferation dynamics of various CSCs upon glucose stimulation. The steady-state level (in molecules) of stem cell marker **(A)** Nanog and **(B)** differentiation marker protein (TGD) are plotted as a function of glucose for BTICs. SN_1_ and SN_2_ represent the saddle node points at 5.1 mM and 2.62 mM glucose, respectively. The steady state levels of **(C)** Nanog and **(D)** differentiation protein (TGD) (in molecule) as a function of glucose in PANC-1, GS-Y03, and A2780 cells. The corresponding glucose level at SN_1_, SN_2_, SN_3_, and SN_4_ (saddle nodes) is 4.9 mM, 3.3 mM, 10.95, and 10 mM. The red lines represent a stable steady state whereas black lines indicate unstable steady states.

This brings the next important question, can the same network model (**Fig. 1**) be used to reconcile the proliferation pattern of CSCs originated from pancreatic cancer (PANC-1), glioblastoma (GS-Y03), and ovarian (A2780) cancer? To unravel this, we did an extensive literature survey and found that in the PANC-1 cell-type, the active STAT3 (STAT3_a_) level remains much higher compared to the active p53 (p53_a_) level (Chen et al., 2018). We must mention that in the case of BTICs, we assume that the STAT3 and p53 expressions remain at a moderate level based on experimental literature.

Thus, we considered a higher expression of STAT3 (by increasing total STAT3 amount) in comparison to p53 level for PANC-1 cell-type (to make sure active STAT3 is higher in amount than active p53 level) and performed our bifurcation analysis keeping everything else the same. The steady state of Nanog shows a reverse mushroom kind of bifurcation as a function of increasing glucose concentration. According to this bifurcation though one can explain the experimentally observed proliferation and differentiation trend of PANC-1 CSCs. The expression level of stem cell marker (Nanog protein) remains in a lower (**Fig. 2C**) steady state and the differentiation protein (TGD) is at a much higher level (**Fig. 2D**) at low glucose concentration (5 mM glucose). However, Nanog reaches a higher steady state in presence of high glucose concentration (26.2 mM glucose) as reported by Shibuya et al. (Shibuya et al., 2015).

Although we did not find any information about the relative levels of active STAT3 and p53 for GS-203, and A2780 cell-types, but our analysis predicts that it could follow a similar trend like PANC-1 cell-type, where the second bi-stable switch of the reverse mushroom bifurcation allows us to explain the proliferation and differentiation pattern under low or high glucose concentrations. Whether a bi-stable switch underlies the transition from proliferation to differentiation in either type of CSCs needs further experimental verification. However, our analysis provides one way to explain the disparate proliferation dynamics of CSCs in terms of bi-modal Nanog regulation. A bi-modal Nanog dynamics will make the cellular decision-making much more robust nonetheless and there is evidence of similar regulation in mammalian cells in the context of proliferation maintenance. Our model further predicts that for cell-types like PANC-1, the stem cell markers (like Nanog) can remain in a high state at very low glucose concentrations (lower than 2.5 mM glucose) which also demands experimental confirmation. We will revisit this issue in the latter part of our discussion.

### p53 preferentially regulates Nanog to maintain differentiated state of BTICs at higher glucose doses

In the previous section, we showed that bi-stable Nanog dynamics in a context-dependent manner qualitatively elucidates the proliferation propensities in CSCs. What molecular mechanisms are governing this disparate proliferation regulation, however, was not clear. Further investigations are due to decipher the key molecular regulators behind these observations. In this regard, we found that overexpressing STAT3 causes an increase in the proliferation of BTICs experimentally (Leidgens et al., 2017). Can our model show a similar response under STAT3 overexpression? We immediately tested it out by considering higher total STAT3 expression compared to WT (**Fig. 3A**). The bifurcation analysis under this condition revealed that the steady state level of Nanog as a function of glucose initially changed from a bi-stable to a reverse mushroom kind of bifurcation (**Fig. 3A, the situation I**), as the STAT3 level was moderately elevated. However, further overexpression of STAT3 (3-fold compared to WT) caused a transition to a higher expression level of Nanog even at higher glucose concentration (**Fig. 3A, situation II**). This implies that if one starts the experiment with very low glucose concentration and keeps adding more glucose in the medium, there will not be any switching between Nanog high to low state with increasing glucose level like in the case of WT BTICs. Thus, our modeling analysis confirms that BTICs have a prominent tendency to go for proliferation with STAT3 overexpression.

**Fig. 3.**
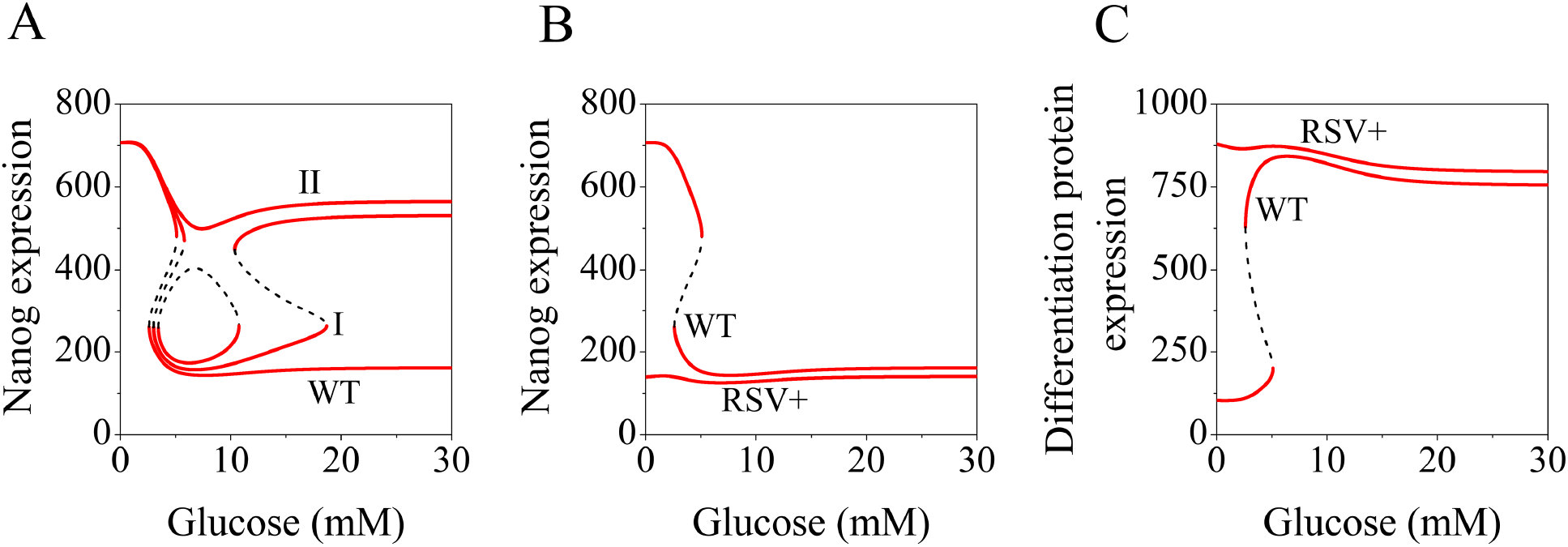
The effect of p53 supersedes STAT3 activation to regulate proliferation dynamics in BTICs. **(A)** Nanog steady state dynamics are plotted concerning glucose concentration for different total STAT3 levels (**I** = 2-fold increase, **II** = 3-fold increase). Steady state levels of **(B)** Nanog and **(C)** differentiation protein (TGD) (in molecules) are plotted against glucose concentration in presence of resveratrol.

Importantly, this result indicates that, to begin with, in the WT scenario for BTICs, the effect of glucose-mediated negative regulation via p53 over Nanog dictates the overall Nanog dynamics by countering the STAT3 facilitated positive activation of Nanog. However, we need a bit more conclusive evidence than this to confirm it. To investigate this aspect, we added the effect of Resveratrol (RSV, trans-3,4’,5-trihydroxystilbene) in our model. RSV is naturally found in grapes, vegetables, and berries, and frequently gets used in various cancer treatments (Heo et al., 2018; Jang et al., 1997). Under normal circumstances, RSV activates p53 by producing ROS (Seino et al., 2015). Furthermore, RSV is known to enhance the differentiation of glioblastoma via ROS dependent signaling pathway (Yuan et al., 2012). We introduced this RSV regulation in our model again in a phenomenological manner (**Table-S1**) and performed the bifurcation analysis for the BTICs in presence of 10 μM RSV (as considered in the experiment) (Yuan et al., 2012). Under such a condition, the Nanog steady state dynamics remain consistently low at any glucose concentration (**Fig. 3B**), while the differentiation related protein shows a steady high expression level indicating the increasing chances of BTICs opting for the differentiated state (**Fig. 3C**). This signifies that RSV further enhances the stronghold of p53 over Nanog dynamics in comparison to STAT3 in BTICs.

### EGF plays a significant role in the proliferation of cancer stem cells

The analysis made in the previous section further suggests that for CSCs originating from PANC-1 kind of cell-types, the STAT3 mediated activation of Nanog might play a significant role in managing the proliferation of CSCs. Thus, the influence of the EGF regulation on Nanog will also become quite important. In this context, Han et al. showed that the proliferation of PANC-1 cells can be modulated by controlling the EGF signaling. In presence of an EGF neutralizing antibody, the proliferation rate decreases, and the threshold of glucose concentration for which proliferation sets (SN_3_ saddle node) becomes much higher than WT. It signifies that glucose not only activates the stem cell markers like Nanog via EGF, but there is another pathway independent of EGF, which is directly regulated by glucose. To understand this phenomenon of PANC-1 cells, we have performed bifurcation analysis as a function of glucose both in the absence (**Fig. 4A-B**, WT) and in the presence (**Fig. 4A-B**, **EGF−**) of EGF neutralizing antibody. The bifurcation analysis in absence of EGF (**Fig. 4A-B**, **EGF−**) displays that a higher threshold level of glucose is required for active proliferation of CSCs, wherein the EGF level shows a significant increase as the glucose level changes from 5.5 mM and 25 mM glucose (**Fig. 4B**) as observed by Han et al. (Han et al., 2011). Assuming that the rate of proliferation is directly proportional to Nanog expression level in stem cells (Chen et al., 2012; Gawlik-Rzemieniewska and Bednarek, 2016), we can say that there is a decrease in proliferation rate in PANC-1 cells (**Fig. 4D**) in absence of EGF as inferred by Han et al. experimentally.

**Fig. 4.**
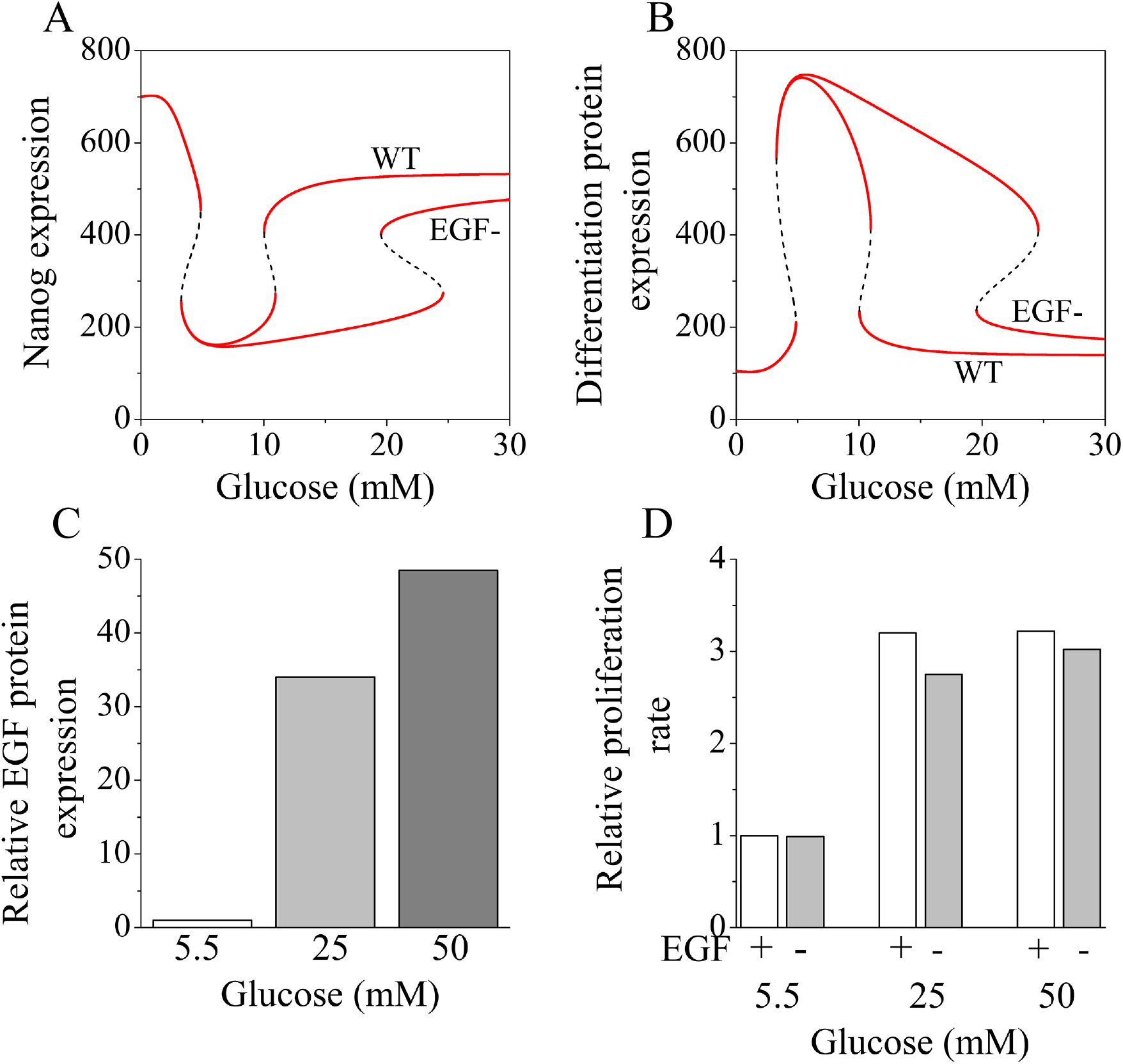
Role of EGF signaling in CSCs proliferation. The variation of steady state **(A)** Nanog and **(B)** differentiation protein dynamics are depicted in absence of EGF in comparison to WT for PANC-1 CSCs. **(C)** The expression of EGF (in molecules) is measured at different levels of glucose concentration. The relative expression levels of EGF are measured from the model with respect to the EGF concentration at 5.5 mM glucose concentration. **(D)** The proliferation rates are plotted for different doses of glucose in the presence and absence of EGF. The proliferation rate is considered to be directly proportional to the Nanog expression. The rate is normalized with respect to the Nanog expression at 5.5 mM glucose in presence of EGF.

To understand the contribution of EGF in the proliferation of CSCs, we blocked the direct activation of STAT3 by glucose and also blocked the activation of STAT3 via EGF, separately. One can see that in absence of direct STAT3 activation, the cells need a little higher concentration of glucose compared to the wild type (WT) to activate the proliferation of CSCs, but there is not much change in the Nanog expression level once it reaches the ON state (**SFig. 1A, II**). Thus, the direct STAT3 activation has a little contribution to stem cell proliferation maintenance, whereas the absence of EGF affects both the threshold glucose level and Nanog expression substantially for CSCs of specific origin under consideration.

Therefore, STAT3 activation via EGF is needed for maintaining the proliferation of PANC-1 cells. However, there is not much change in the proliferation dynamics of BTICs in the presence or absence of EGF (**SFig. 1B**). This elucidates that the negative regulation by p53 concerning Nanog activation acts in a dominating manner and counters the effect of EGF and STAT3 mediated Nanog activation to maintain the proliferation response in BTICs.

### Resveratrol induced loss of self-renewal in CSCs via ROS mediated p53 activation

However, it seems that for cell-types like PANC-1, this happens in an opposite manner to decide the proliferation outcome of CSCs. To further illustrate this matter, we concentrate on this section to understand the effect of p53 mediated Nanog regulation in these CSCs. We have already discussed earlier that RSV treatment in a specific way can allow us to decipher the extent of the p53 effect over Nanog. This is because RSV alters the ROS activation and in turn affects p53 regulation in these CSCs in general. In this regard, Seino et al. (Seino et al., 2015) had shown that in A2780 cancer stem cells, resveratrol treatment induced differentiation of CSCs via ROS activation. Can our model reproduce the experimental observations made by Seino et al.? To answer this question, we have performed our modeling analysis at 26.2 mM glucose concentration and in presence of 50μM RSV. We found that the ROS level increased five-fold (**Fig. 5A**), whereas the stem cell markers (Nanog and Sox2) decreased significantly compared to control (without RSV) (**Fig. 5B**). Following the experimental protocol, next, we calculated the ROS level and expression of stem cell markers in presence of N-acetylcysteine (NAC) (20 nM). NAC treatment is known to effectively reduce the ROS level and inhibit DNA damage (Xie et al., 2018). We observed that upon treatment with NAC, the ROS level reduced to less than half the amount compared to the control (**Fig. 5A**), as observed in the experiments with A2780 cells, while the stem cell markers increased marginally (**Fig. 5B**). Following the experiments, introducing NAC along with RSV is neutralizing the effect of RSV mediated ROS induction in presence of 26.2 mM glucose (**Fig. 5A-B**).

**Fig. 5.**
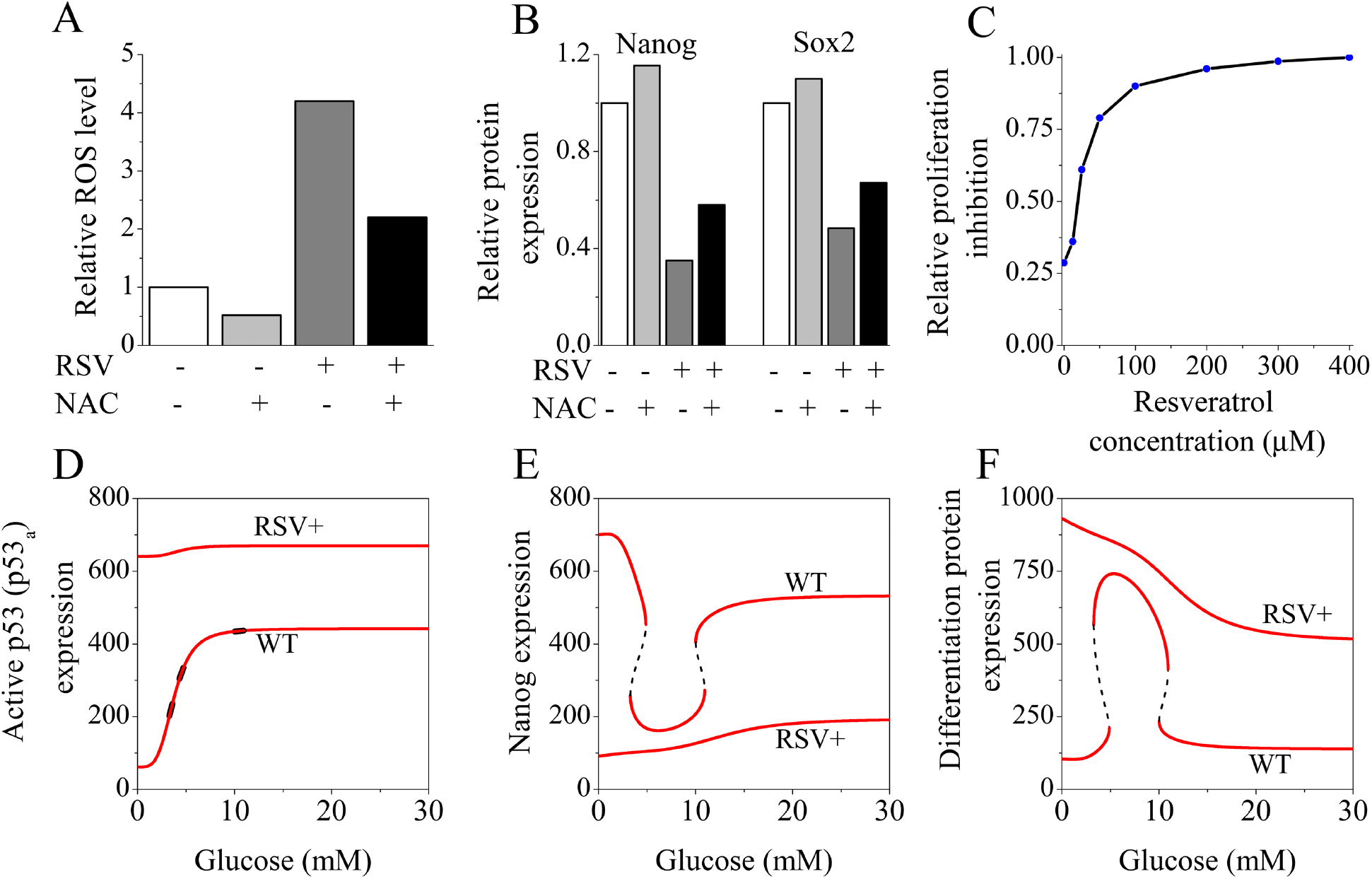
The role of resveratrol in controlling the cellular proliferation of CSCs via ROS and p53 mediated Nanog regulation. **(A)** The relative level of ROS is calculated at 26.2 mM glucose concentration from the model under different experimental conditions by normalizing them to the WT ROS level. **(B)** The model predicted expression levels of stem cell markers (Nanog and Sox2) are shown in presence of both RSV (50 μM) and NAC (20 nM) separately and presence of a mixture of both RSV (50 μM) and NAC (20 nM). **(C)** The relative proliferation inhibition is shown as a function of increasing doses of Resveratrol (we considered that proliferation inhibition rate is proportional to the inverse of the Nanog concentration and it is normalized to the Nanog expression in presence of 400 μM RSV). We have plotted the steady state levels of **(D)** active p53 (p53_a_), **(E)** Nanog, and **(F)** differentiation protein (TGD) in the presence (RSV+) and absence (WT) of resveratrol.

Other experimental studies in similar lines had shown the anticancer effect of RSV on pancreatic carcinoma cells (Cui et al., 2010; Harikumar et al., 2010; Oi et al., 2010). In this context, Mo et al. (Mo et al., 2011) had measured the percentage inhibition of cell proliferation by RSV compare to the control group and concluded that resveratrol inhibited the growth of PANC-1 cells in a dose-dependent manner. We have implemented a similar modeling calculation of the inhibition of proliferation (assuming that the inhibition of proliferation is inverse of Nanog expression) with respect to varying concentrations of RSV (**Fig. 5C**) by calculating the proliferation rate for a reasonably higher glucose concentration (50 mM). The choice of higher glucose concentration was made because at high glucose levels the respective CSCs must be highly proliferative according to the bifurcation diagram (**Fig. 2C-D**). Our result displays a direct relationship between RSV and cell proliferation inhibition (**Fig. 5C**) as shown by Mo et al. (Mo et al., 2011). Is it happening by restricting p53 activation?

p53 activation is an important factor in regulating Nanog activation within CSCs, and treatment with resveratrol was also found to activate the p53 pathway (Athar et al., 2009; Sato et al., 2013). In this context, Sato et al. had shown that resveratrol promotes differentiation and suppresses the self-renewal capacity of GS-Y03 cells. They also have demonstrated that resveratrol induces the phosphorylation of p53 to an active form of p53 (p53_a_), which is the major reason for the suppression of Nanog. We performed a bifurcation study of similar nature and found a substantial increase in the expression of active p53 in presence of 50μM RSV under any specific glucose concentration (**Fig. 5D**). Importantly, upon RSV treatment, the bi-stable dynamics of Nanog vanishes (**Fig. 5E**), and Nanog eventually maintains a very steady low level throughout as a function of varying glucose concentration. This demonstrates the onset of the differentiation program of CSCs and it even reflects in the dynamics of the differentiation related protein (**Fig. 5F**) as well. In GS-Y03 cells, overall STAT3 expression is much higher than the p53 expression, however, in presence of RSV, p53 gets highly activated through ROS and overcomes the influence of STAT3 on Nanog. Once active p53 takes over the proceedings, it destroys the positive feedback regulation on Nanog, hence Nanog loses its bi-stable dynamics. This result signifies that even in GS-Y03 cells, the proliferation inhibition through Nanog response is mainly governed by the activation of p53 via ROS activation.

### Sensitivity analysis of model parameters reveals ways to fine-tune the proliferation dynamics of CSCs

Till now, our model has shown promise by reproducing several experimental phenomena related to proliferation tendencies of CSCs. Can it further predict how to fine-tune the proliferation propensities of CSCs in a context-dependent manner? To address this question, we performed a systematic sensitivity analysis (**SFig. 2**) of the model parameters (**Table-S3**) by carefully opting for the positions of bifurcation points (**Fig. 2**) as the sensitivity parameter to explore the possible ways to influence the cell-type dependent proliferation CSCs in terms of Nanog dynamics. Here, we intended to identify the model parameters, by altering which we can tweak the proliferation/differentiation response in a cell type-specific manner. To begin with, we consider the case of BTIC cell-type. In **Fig. 2A**, it was evident that the positions of the saddle nodes SN_1_ and SN_2_ critically control the threshold of glucose concentration beyond which the cell will start to differentiate leaving the proliferation program.

We picked up a specific example of p53 related parameters from **SFig. 2** to depict the strength of our analysis. **Fig. 6A-B** demonstrates that increasing the amount of total p53 will push both the saddle nodes (SN_1_ and SN_2_) towards the lower level of glucose. This suggests that with increasing total p53 level, BTICs will require less glucose concentration to opt for differentiation (**Fig. 6C**). However, a further decrease in the total p53 level will allow the cells to proliferate unconditionally, as it maintains Nanog completely in the ON state (**Fig. 6C, I**). This indicates that the cancer cells (BTICs) will become even more aggressive when any parameter associated with the production of p53 is downregulated. However, only ROS activation can reverse the scenario (**Fig. 6A-B**) as it pushes both the saddle nodes towards a higher glucose threshold. This exhibits that by only altering the p53 dynamics, one can even drive the BTICs towards either proliferation or differentiation depending on the demand of the situation.

**Fig. 6.**
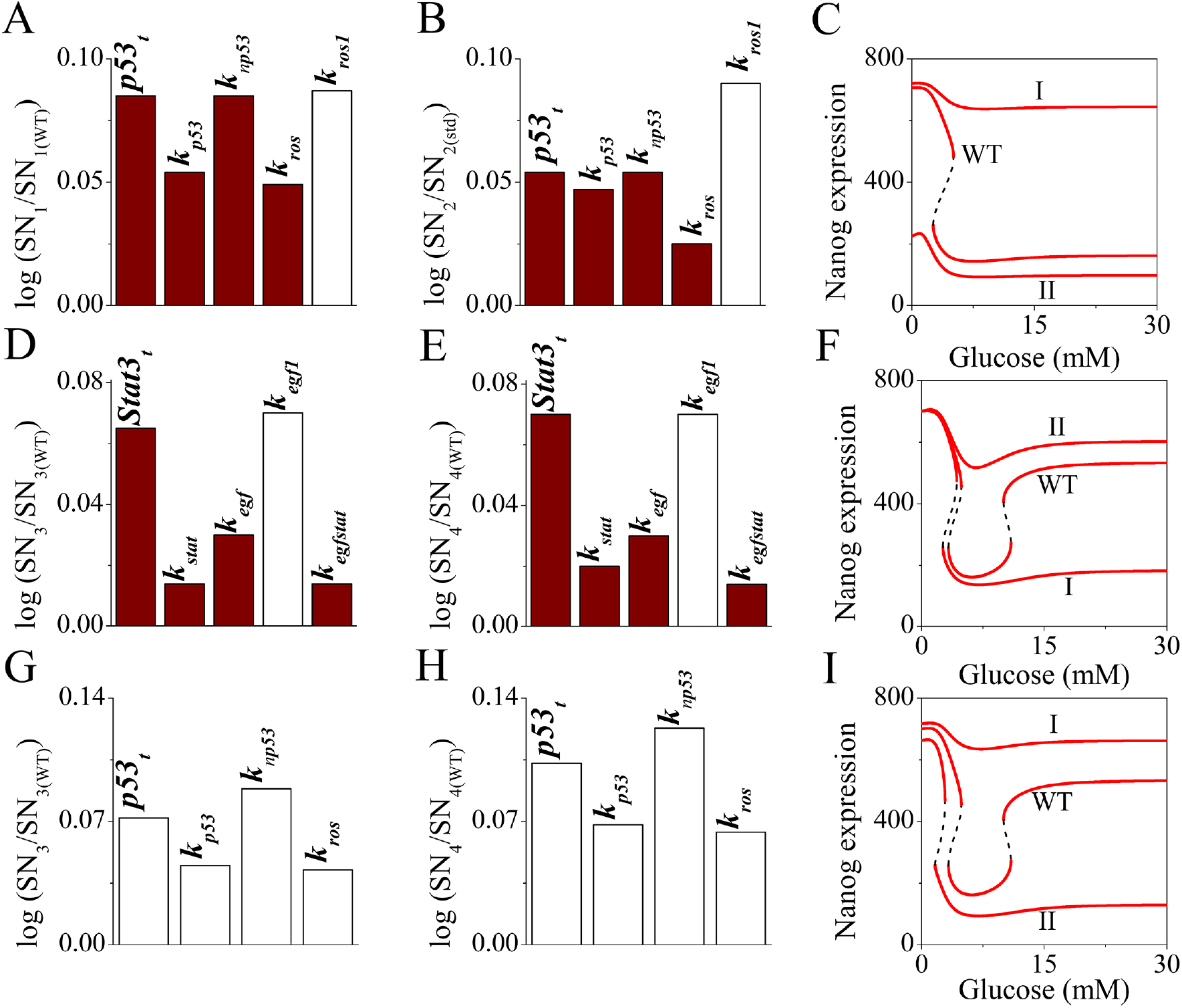
Sensitivity analysis reveals the effect of glucose-mediated down-stream regulators on the proliferation of CSCs. The white and brown bars represent positive and negative regulation respectively. The sensitivity analysis of the parameters related to p53 is plotted by considering **(A)** SN_1_ saddle node position and **(B)** SN_2_ saddle node position (**Fig. 2A**) related to BTICs. **(C)** The steady-state Nanog dynamics is plotted as a function of glucose for a different level (I (p53_t_ level is 0.5 times of WT) and II (p53_t_ level is 2 times of WT)) of total p53 (p53_t_) in comparison to WT case. The sensitivity analysis of the parameters related to STAT3 is plotted by considering **(D)** SN_3_ saddle node position and **(E)** SN_4_ saddle node position (**Fig. 2C**) related to PANC-1 and related cell-line CSCs. **(F)** The steady state dynamics of Nanog as a function of glucose are shown for varying concentrations of total STAT3 (I (STAT3_t_ level is 0.5 times of WT) and II (STAT3_t_ level is 2 times of WT. The sensitivity analysis of the parameters related to p53 total (p53_t_) are plotted by considering **(G)** SN_3_ saddle node position and **(H)** SN_4_ saddle node position (**Fig.2C**) related to PANC-1 and related cell-line CSCs. **(I)** The steady state dynamics of Nanog as a function of glucose are shown for varying concentrations of total p53 (I (p53_t_ level is 0.5 times of WT) and II (p53_t_ level is 2 times of WT)).

On the other hand, for the PANC-1, A2780, and GS-Y03 CSCs scenario, we have mainly focused on the saddle nodes SN_3_ and SN_4_ to perform the sensitivity analysis (**SFig. 3** and **Fig. 6D-I**), as we already have mentioned that there is no experimental report of following Nanog dynamics below 5 mM glucose concentration for these CSCs. Interestingly, all the parameters that appear in **Fig. 6D-I**, in one way or the other are related to glucose activated upstream regulators (p53 and STAT3) of Nanog dynamics. Our sensitivity calculations predict that for CSCs of PANC-1, A2780, and GS-Y03 origin, the proliferation tendency will increase with an elevated level of STAT3 (**Fig. 6D-E**). This is because an increase in total STAT3 level maintains the CSCs at the proliferative state under any glucose concentration (**Fig. 6F, II**). However, the cells will remain differentiated if STAT3 gets inhibited (**Fig. 6F, I**) for a fixed level of glucose concentration (more than 5 mM). Experimentally, this prediction can be tested by employing certain DNA-alkylating platinum complexes, which induce the death of tumor cells by direct inhibition of active STAT3 (Turkson et al., 2004).

In this regard, EGF regulation will also play a similar important role in cancer stem cell therapy. For this type of CSCs, the parameters related to the activation of EGF shift the respective saddle points towards lower glucose levels (**SFig. 4**). This suggests a higher level of EGF will promote a higher degree of proliferation even in presence of lower glucose concentration. Therefore, instead of knocking down STAT3, one can get the same kind of favorable differentiation of CSCs under consideration by using EGF inhibitors shown earlier in **Fig. 4**. Additionally, by changing the expression level of total p53, one can influence the proliferation dynamics of PANC-1, GS-Y03, and A2780 CSCs (**Fig. 6G-I**). Our analysis depicts that increasing the level of total p53_t_ will induce differentiation of the CSCs, as it suppresses the Nanog expression (**Fig. 6I, II**). Overall, our sensitivity analysis opens up new experimentally testable hypotheses to control the differentiation or proliferation dynamics of cancer stem cells. One way to get rid of these CSCs is to simply create an experimental condition to let them differentiate into a specific cell-state, where they cannot proliferate indefinitely. Recent studies indicate that therapies based on this idea are getting popular to target cancer stem cells (Jin et al., 2017). In this context, our model predictions could be useful to target origin-specific CSCs in a cell-type-specific manner.

## Discussion

Cancer stem cells are highly dependent on glycolysis, fatty acid synthesis, and glutamine metabolism for nutrient consumption. The ‘Warburg effect’ signifies that cancer stem cells favor glycolytic breakdown of glucose for energy intake, although the amount of energy (ATP) produces in mitochondrial oxidative phosphorylation is much higher compare to glycolysis (Chen et al., 2007; DeBerardinis et al., 2008; Gatenby and Gillies, 2007). Thus, to maintain CSCs proliferation, it requires excess glucose to compensate for the insufficient metabolic process of energy production (Snyder et al., 2018). However, in some specific CSCs, it was observed that increasing glucose level leads to differentiation of the corresponding CSCs. The underlying biological network that functions to govern the glucose facilitated proliferation in CSCs is highly complicated and demands a better understanding of the problem, as knowing this regulation properly can open up various therapeutic avenues in the future. Herein, by proposing a qualitative mathematical model of glucose mediate Nanog regulation in cancer stem cells (**Fig. 1**), we investigated the reason behind the disparate proliferation pattern of CSCs under the influence of increasing doses of glucose.

With the proposed model (**Table-S1**) based on **Fig. 1**, we performed a thorough bifurcation analysis (**Fig. 2**), which qualitatively described the experimentally observed differential dynamics of Nanog as a function of glucose for CSCs of different origin. It is important to note here that we only assume the CSCs of various origins have a different ratio of STAT3 to p53 expression levels and found that only this assumption is enough to reconcile the experimentally observed proliferation behavior of CSCs of diverse cancer cell-types. Our analysis suggests that in BTICs, p53 activity dominates over STAT3 regulation (**Fig. 3**) and generates a specific proliferation pattern (**Fig. 2A-B**). However, in pancreatic, ovarian, and glioblastoma CSCs, STAT3 plays a major role in the assistance of EGF signaling (**Fig. 4**). It turns out that for these CSCs, one can further tune the proliferation response by modifying the p53 response employing external agents like Resveratrol or NAC (**Fig. 5**). Essentially our bifurcation analysis (**Fig. 2**) proposes that the proliferation propensities in CSCs are governed by altering the bi-stable Nanog existing within CSCs by administrating different doses of glucose. However, further experimental verifications are due to understand whether glucose controls the bi-stable dynamics of Nanog or not.

In this regard, we found an unusual kind of bifurcation (reverse mushroom, **Fig. 2C**) of Nanog steady state dynamics as a function of glucose, which explains the proliferation observation at 5 mM and 25 mM glucose concentrations for the respective CSCs. However, it indicates that for CSCs that originated from cell-types like PANC-1, A2780, and GS-Y03, there is a possibility that for very less glucose level (say less than 2.5 mM), these cells will transit to a proliferative state due to higher levels of Nanog (**Fig. 2C**). This has not yet been seen experimentally, as none of the experimental work used such a low level of glucose for these cell-types. However, we went one step ahead and tried to figure out if the dynamics of Nanog can be modified further to get rid of this phenomenon of early transitions from proliferation to differentiation for these CSCs. We slightly modified the glucose-mediated ROS activation term in Eq. 7 (**Table-S1**) and few related parameters values (**Table-S3**) to avoid the initial switching of Nanog from a proliferative to a differentiated state below 5 mM glucose concentration by considering the same gene regulatory network (**Fig. 7C-D**). The bifurcation analysis of the model with the modified equation and parameters reproduced similar behavior for BTICs as well (**Fig. 7A-B**). All other modeling results therein remain similar as presented in **SFig. 5-8**. Properly planned experiments in this direction will establish, out of the two possible modeling scenarios, which one more reliably explains the behavior of CSCs of different origin. For both the model types, we have performed a systematic sensitivity analysis (**Fig. 6, SFig. 2-3** and **SFig. 9-11**) to figure out how to fine-tune the proliferation response in different CSCs. The sensitivity analysis predicts that by elevating or down-regulating the key cellular components like p53 and STAT3, how one can regulate the extent of proliferation and differentiation of CSCs in a context-dependent manner.

**Fig. 7.**
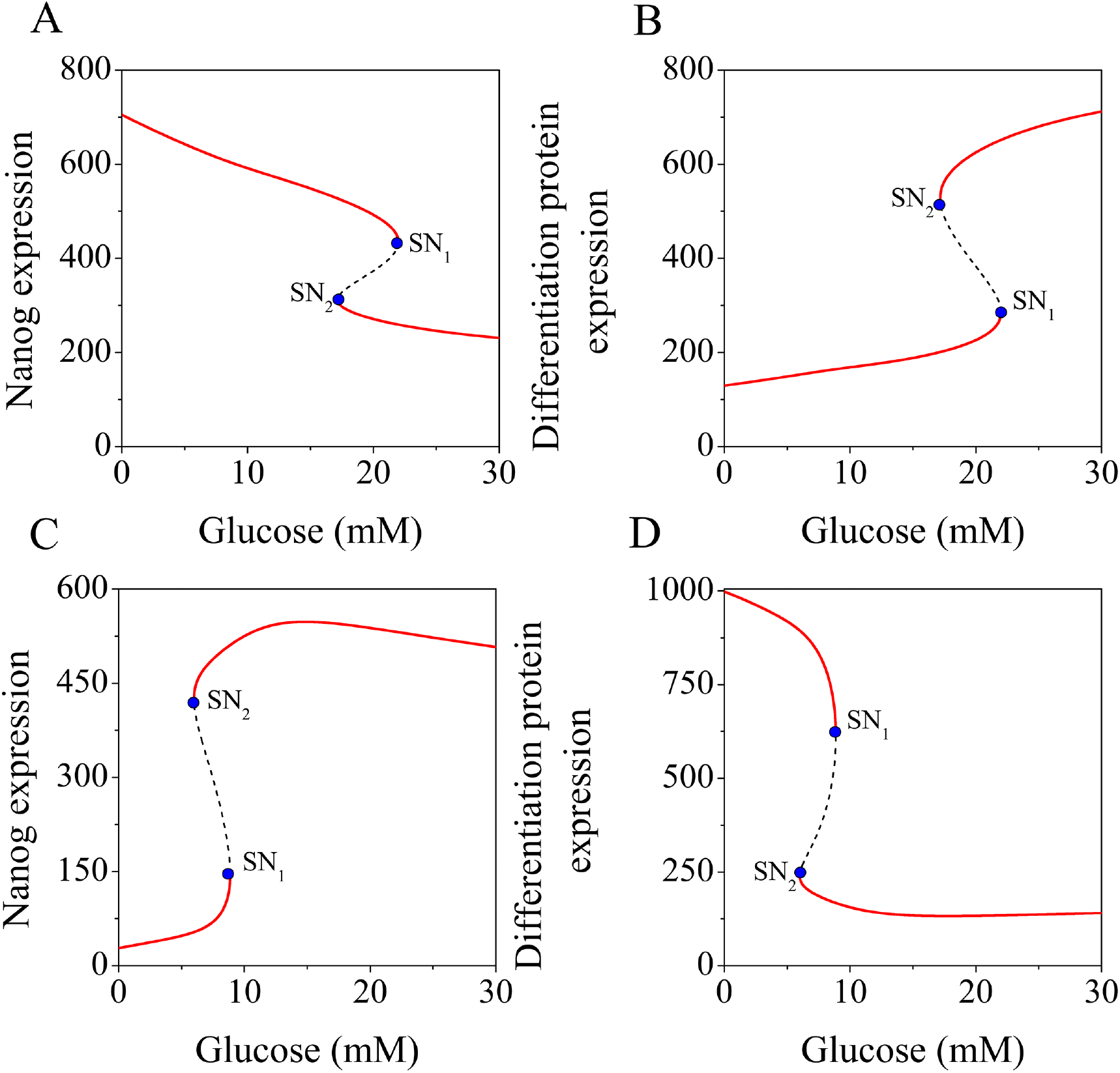
Bifurcation analysis of the model demonstrates the disparate proliferation dynamics of various CSCs upon glucose stimulation. The steady-state level (in molecules) of stem cell marker **(A)** Nanog and **(B)** differentiation marker protein (TGD) are plotted as a function of glucose for the BTIC scenario. The steady state levels of **(C)** Nanog and **(D)** differentiation protein (TGD) (in molecule) as a function of glucose in PANC-1, GS-Y03, and A2780 cells. The corresponding glucose level at SN_1_ and SN_2_ is 12.24 mM and 8.68 mM respectively. The red lines represent a stable steady state whereas black dotted lines indicate unstable steady states.

In conclusion, our proposed model effectively elucidates the glucose-mediated steady-state dynamics of stem cell and differentiation markers in CSCs in a cell-type dependent manner. It foretells qualitatively how one can manipulate the STAT3 or p53 dependent part of our proposed Nanog regulatory network (**Fig. 1**) to direct a particular type of CSC to commit for either differentiation or proliferation. Thus, our modeling study predicts new possibilities to control the unwanted proliferation of CSCs in an improved mode and will be helpful in providing insights to develop therapeutic strategies to eradicate CSCs.

## Method

The proposed gene regulatory network consists of 9 ordinary differential equations. The deterministic model was encoded as an .ode file, and the bifurcation analysis was carried out using freely available software XPP-AUT. The AUTO facility available in XPP-AUT software was used to perform the deterministic bifurcation analysis. The bifurcation diagrams were drawn with the help of OriginLab using data points generated by XPP-AUT.

For the systematic sensitivity analysis, parameters were increased individually at an amount of 20% of the model parameters, keeping all other parameters constant. Sensitivity analysis of the model parameters was performed by considering the concentration of glucose at different saddle points as the sensitivity criteria. White-colored bar signifies an increase in the required concentration of glucose with respect WT case whereas the brown-colored bar signifies a decrease in the expression level of glucose with respect to WT.

## Acknowledgments

Thanks are due to IRCC, IIT Bombay for providing fellowship to (TS). This work is supported by the funding agency SERB, India grant no - CRG/2019/002640.

